# A Dual-Fluorescence Assay for Gene Delivery Vehicle Screening in Macrophages with an Inflammation-Inducible Reporter Construct

**DOI:** 10.1101/2024.08.05.606664

**Authors:** Allie Ivy, Shelby N. Bess, Shilpi Agrawal, Varun Kochar, Abbey L. Stokes, Timothy J. Muldoon, Christopher E. Nelson

## Abstract

**Background:** Macrophages are a promising target for therapeutics in various applications such as regenerative medicine and immunotherapy for cancer. Due to their plastic nature, macrophages can switch from a non-activated state to activated with the smallest environmental change. For macrophages to be effective in their respective applications, screening for phenotypic changes is necessary to elucidate the cell response to different delivery vehicles, vaccines, small molecules, and other stimuli.

**Methods:** We created a sensitive and dynamic high-throughput screening method for macrophages based on the activation of NF-κB. For this reporter, we placed an mRFP1 fluorescence gene under the control of an inflammatory promoter, which recruits NF-κB response elements to promote expression during the inflammatory response in macrophages. We characterized the inflammatory reporter based on key markers of an inflammatory response in macrophages including TNF-α cytokine release and immunostaining for inflammatory and non-inflammatory cell surface markers. We compared gene delivery and inflammation of several clinically relevant viral vehicles and commercially available non-viral vehicles. Statistical analysis between groups was performed with a one-way ANOVA with post-hoc Tukey’s test.

**Results:** The reporter macrophages demonstrated a dynamic range after LPS stimulation with an EC50 of 0.61 ng/mL that was highly predictive of TNF-α release. Flow cytometry revealed heterogeneity between groups but confirmed population level shifts in pro-inflammatory markers. Finally, we demonstrated utility of the reporter by showing divergent effects with various leading gene delivery vehicles.

**Discussion:** This screening technique developed here provides a dynamic, high-throughput screening technique for determining inflammatory response by mouse macrophages to specific stimuli. The method presented here provides insight into the inflammatory response in mouse macrophages to different viral and non-viral gene delivery methods and provides a tool for high-throughput screening of novel vehicles.

## Introduction

Macrophages are phagocytic cells responsible for defense against foreign invaders and maintaining homeostasis in all organs and tissues^1–3^. Based on the microenvironment, macrophages change function to respond to local need. The plasticity of macrophages leads to a heterogenous population of macrophage phenotypes to address the situation, whether defense, maintenance, or in transition between activation states. Macrophages play a role in tumors as tumor-associated macrophages (TAMS) and regenerative processes in the body. For many cancers, macrophages are abundant in the tumor microenvironment and TAMS are responsible for facilitating metastasis, immunosuppression, and promoting invasion and angiogenesis^4^. Macrophages are also responsible for maintaining the healing process from initial inflammation to remove foreign invaders, recruiting necessary immune cells, and resolving the healing process in the end stages of regeneration^5–9^.

Macrophages can participate in a wide variety of activities due to their ability to switch between activation states. Understanding of macrophage polarization state is continually evolving and at the most basic level is either a classically activated/inflammatory or alternatively activated/anti-inflammatory state. These states have also been described as M0 (resting), M1 (inflammatory), and M2 (anti-inflammatory). Due to their utility, macrophages have been targeted for use in many different applications from cell therapy for oncology to reprogramming the local environment in regeneration^10–16^. Although macrophages provide a range of utility, their ability to switch between an activated and alternatively activated state provides challenges in adapting these cells for use in different therapeutic applications. With the advent of genetically engineered macrophage cell therapy^16^, the role of delivering gene-editing machinery to macrophages has been of increasing interest as viral and non-viral delivery methods can prove to be highly inflammatory.

The landscape of different gene-editing tools available for use has dramatically changed over the past few decades. These new tools and techniques provide a way to both investigate and manipulate cellular behavior and are invaluable in the development of therapeutics targeting and using macrophages. So far, macrophage gene expression has been manipulated using shRNA^17^, siRNA^18–20^, zinc-fingers (ZFNs)^21^, transcriptional activator-like effector nucleases (TALENs)^22,23^, and CRISPR-Cas9^24–26^. And yet, delivery of gene-editing tools remains a challenge. Due to the innate nature of macrophages to respond to foreign invaders, many viral and non-viral delivery vehicles lead to an inflammatory response^16,27^. Although this may be desirable for some approaches, as mentioned before, many therapeutic applications that would benefit from macrophage manipulation require an anti-inflammatory phenotype favoring regenerative processes, such as for chronic wounds^9^, skeletal muscle damage following injury^28^, myocardial infarction^29–31^, impaired liver regeneration^32^, and even brain damage^33,34^.

For cancer immunotherapy, inflammatory macrophages are beneficial as that is the desired state for the cell therapeutic, but for regenerative approaches, an inflammatory macrophage phenotype would prove detrimental to the continued healing process. It is essential to determine the effects of different delivery modalities on macrophages before attempting to manipulate them with current gene-editing technology.

To screen the inflammatory response of macrophages to different delivery vehicles, we developed a high-throughput screening method that relays the inflammatory response of RAW264.7 mouse macrophage cells to viral and non-viral delivery methods. Others have taken similar approaches to investigate macrophage inflammatory responses using nuclear factor kappa B (NF-κB) reporters, an important transcription factor for the inflammatory response, to determine which genes regulate inflammatory pathways in macrophages^35^. Here, we establish a high-throughput screening technique to determine macrophage inflammatory response via the NF-κB activation pathway. We then used the high-throughput screen for viral and non-viral delivery vehicles to determine their effect on macrophage activation.

## Methods

### Cell lines

The RAW264.7 murine macrophage cell line was purchased from the American Type Culture Collection (ATCC TIB-71). These cells were cultured in Dulbecco’s modified Eagle’s medium (DMEM) (Gibco) with 10% fetal bovine serum (FBS) (Gibco) and 1% penicillin/streptomycin (P/S) (Gibco). Cells were grown in a tissue culture incubator at 37°C and 5% CO_2_. Cells were passaged using a cell scraping method to resuspend the adherent cells.

### Plasmid construction

The inflammatory reporter plasmid was constructed with mRFP1 red fluorescent protein under a previously described synthesis-friendly inflammation-inducible promoter (SFNp) with puromycin resistance under a PGK promoter and lentivirus packaging elements. The plasmid pLenti-CMV-GFP-Puro (Addgene #17448) containing lenti packaging elements, GFP under a CMV promoter and puromycin resistance under a PGK promoter, was used as the backbone. The CMV-GFP was removed from the plasmid using ClaI and SaII and purified via gel electrophoresis. The mRFP1 red fluorescent protein was taken from the pU6-pegRNA-GG-acceptor plasmid (Addgene #132777) via PCR using Q5 High-Fidelity DNA Polymerase (NEB) and purified. The SFNp sequence was ordered as a gene block from Integrated DNA Technologies (IDT). The SFNp sequence is:

> GGATCCACGGGATACCCCAGGGGCTCTCCAGGGAATCTCCGGGGATACTCCAGGGGGTTTCCGGGGAATCC CCCGGGAGTTTCCTGGGAATTTCCCGGGATTTCCCCGGGGCATCCCGGGGACTCTCCTGGGATTTTCCAGG GACATTCCTGGGACTTTCCTGCGCGGTAGGCGTGTACGGTGGGAGGTCTATATAAGCAGAGCTCGTTTAGT GAACCGTCAGATCGCCTGGAGACGCCATCCACGCTGTTTTGACCTCCATAGAAGACACCGGGACCGATCCA GCCTCTCGACATTCGTGCCACCATG^36^.

Overhangs were added to the gene block via PCR and purified. All PCR primers for overhangs were designed using NEBuilder Assembly Tool. NEB Gibson Assembly® Master Mix was used to combine the fragments according to manufacturer’s instructions.

### Stable cell culture creation using lentivirus

The inflammatory reporter plasmid was used to create a stably expressing cell line via lentiviral infection, using the 2^nd^ generation lentiviral system including psPAX2 (Addgene #12260), pMD2.G (Addgene #12259), and the inflammatory reporter plasmid. HEK293T cells were seeded in 10cm plates at a seeding density of 2 million cells. At 75-90% confluency, all three plasmids were transfected into HEK293 cells using calcium phosphate precipitation transfection. Media was changed 16 hours post-transfection, and media containing lentivirus was harvested at 24 and 48 hours post-media change. Supernatant containing lentivirus was then filtered using a 0.45 µm cellulose acetate filter. Lenti-X concentrator was added to the filtered viral supernatant at a 1:4 ratio and stored at 4°C overnight. The following morning, the centrifuge was cooled to 4°C and the virus was centrifuged at 1,500x*g* for 45 minutes. Supernatant was aspirated and pellet resuspended in PBS at 20x the volume of the original viral media. RAW264.7 cells were seeded in 5cm plates at a seeding density of 800,000 cells. At 75% confluency, lentivirus containing the inflammatory reporter was added. Viral media was replaced with fresh media 24 hours post-transduction. The cells were allowed to expand for another 24 to 48 hours before puromycin selection occurred with a concentration of 3.5 µg/ml for one week. Following puromycin selection, cells were activated using LPS at 1 ng/ml over 24 hours before using flow sorting to obtain the top 10% of mRFP1 expressing cells. Cells were expanded and then cryo-preserved for future use.

### Viral Transduction of RAW264.7-IRCs

RAW264.7-IRCs were seeded in 24-well plates at a seeding density of 50,000 cells per well. At 75% confluency, media was changed 2 hours before transfection. The RAW264.7-IRCs were transduced with AAV 1, 2, 5, 8, and 9, all containing a GFP construct (UNC Vector Core), and Adenovirus 5/35 (Ad5/35)-GFP (Welgen, Inc.). Inflammatory reporter RAW264.7 cells were transduced at an MOI of 38,000 for all AAV serotypes and 1,000 for Ad5/35^16^. Transfection media was replaced with fresh media 24 hours post-treatment and cells were allowed to expand for another 24 or 48 hours, respectively, before harvesting cells for flow cytometry.

### Non-Viral Transfection of RAW264.7-IRCs

RAW264.7-IRCs were seeded in 24-well plates at a seeding density of 50,000 cells per well. At 75% confluency, media was changed 2 hours before transfection. Transfection using Lipofectamine^TM^ 2000 (ThermoFisher) and TransIT-X2® Dynamic Delivery System (Mirus bio), were carried out per manufacturer instructions. Briefly, 5 µg of an expression plasmid containing GFP was diluted in Opti-MEM^TM^ Reduced Serum Medium (Gibco^TM^) and mixed with Lipofectamine 2000 diluted in Opti-MEM. The DNA/Lipofectamine 2000 mixture was incubated for 5 minutes at room temperature before being added to the RAW264.7-IRCs. Cells were incubated for 48 hours before harvesting. For TransIT-X2, it was brought to room temperature and vortexed prior to use. 1.5 µg of pDNA was diluted in 150uL Opti-MEM. 4.5 µl of TransIT-X2 was then added to the diluted DNA mixture and incubated for 15-30 minutes at room temperature before being split evenly across three wells. Cells were incubated for 48 hours before harvesting.

### Flow cytometry

All flow cytometry experiments were performed using the MA900 instrument (Sony Biotechnology). All cells were re-suspended in Dulbecco’s phosphate buffered saline (PBS) (ThermoFisher) via scraping methods prior to sorting. Samples were strained using EASYstrainer Cell Sieves 40 µm (Greiner Bio-One). All stimulated RAW264.7-IRCs were evaluated alongside unstimulated RAW264.7-IRCs and RAW264.7 cells lacking any reporter for initial gating and to set the mRFP1-gate. Cells were gated first to determine live cells by forward (FSC-A) and side scatter (SSC-A) and then FSC-A and FSC-H for single cells. Cells were then evaluated for mRFP1 (SSC v. mRFP1) and/or GFP expression (SSC v. GFP). Cells were discarded after flow cytometry analysis. Further information on the gating strategy can be found in **Supplementary Figure S4**. All data was evaluated using FlowJo™ Software.

### Immunostaining

Immunostaining was performed to characterize phenotypic surface expression post-cytokine stimulation. AlexaFluor™ 647 secondary antibody with CD86 primary antibody was used to detect CD86, a common M1 marker, and AlexaFluor™ 488 secondary antibody with CD206 primary antibody was used to detect CD206, a common M2 marker. The RAW264.7-IRCs were seeded in 35 mm glass bottom MatTek dishes, P35G-1.5-10-C (MatTek Life Sciences) at a seeding density of 300,000 cells per dish, 24 hours prior to cytokine stimulation. LPS at 10 ng/mL was used for stimulation. 24 hours post-cytokine stimulation, cells were fixed in 10% neutral buffered formalin and permeabilized using 0.2% Triton-X 100, followed by blocking with 2% bovine serum albumin (BSA) for 60 minutes. The primary antibodies were added and allowed to incubate overnight at 4°C. Cells were then washed with phosphate buffered saline (PBS) prior to imaging. Images were acquired using an inverted laser scanning confocal microscope (Olympus Fluoview FV10i-LiV) with a 60X (1.2□N.A., water immersion).

### Antibody Staining for Flow Cytometry

The RAW264.7-IRCs were seeded in 60 mm plastic dishes at a seeding density of 1,000,000 cells per dish. RAW264.7 cells were plated to control for mRFP1. The next day, media was either replaced with normal media or media containing LPS (10 ng/mL). 24 hours post stimulation cells were collected and resuspended in 400 μL of PBS containing 1% BSA. The cells from each dish were split in half to be stained for either CD86 or CD206. CD86 and CD206 primary antibody were used at a 1:20 and 1:15 volume dilution respectively and incubated for 1 hr. After incubation, cells were washed with PBS containing 1% BSA three times. The AlexaFluor™ 647 secondary antibody (CD86) and AlexaFluor™ 488 secondary antibody (CD206) were added to a final concentration of 5μg/mL and incubated for 30 minutes. After incubation, cells were washed and resuspended in 300 μL of PBS containing 1% BSA. Cells were subsequently sorted using similar machinery and gating strategies described above in the Flow Cytometry section.

### TNF-α ELISA

RAW264.7-IRCs were seeded in 24-well plates at a seeding density of 50,000 cells/well. 24 hours after seeding, cells were washed with PBS and media were replaced with stimulating media at varying doses of LPS (0.001 ng/mL to 1 ng/mL) for 24 hours. Supernatant was collected and spun down to remove any cellular debris and then frozen at -80°C prior to use. The cell culture supernatant was thawed and diluted 1:10. TNF-α levels were measured using the TNF alpha Mouse Elisa Kit (Invitrogen) according to manufacturer instructions. Absorbance readouts were analyzed using a BioTek Synergy LX multi-mode reader (Agilent), measured at 450nm. The standard curve was graphed and used to determine the concentration of TNFα of each sample. Concentrations were corrected for dilution factor.

## Results

### Inflammatory Reporter Characterization

To achieve high-throughput screening of macrophage’s inflammatory response to external elements, an inflammatory reporter plasmid was constructed. The coding sequence for mRFP1 fluorescent protein placed under an inflammation-inducible promoter from Jadav and Truong^36^ using Gibson cloning. The inflammatory promoter contained 14 NF-κB transcription factors with a minimal CMV promoter. The plasmid also contained lentivirus packaging elements with puromycin resistance under an Ef1α promoter to maintain expression over time (**Fig. 1A**). The plasmid sequence was validated using Sanger sequencing. To evaluate the inflammatory reporter, it was stably integrated in RAW264.7s, a mouse macrophage cell line, using lentiviral transduction^37^. Following puromycin selection, the RAW264.7-Inflammatory reporter cells (RAW264.7-IRCs) were stimulated using lipopolysaccharide (LPS) from *E. coli and* sorted for the top 10% of mRFP1 expressing cells to ensure a more homogenous population.

**Figure 1.**
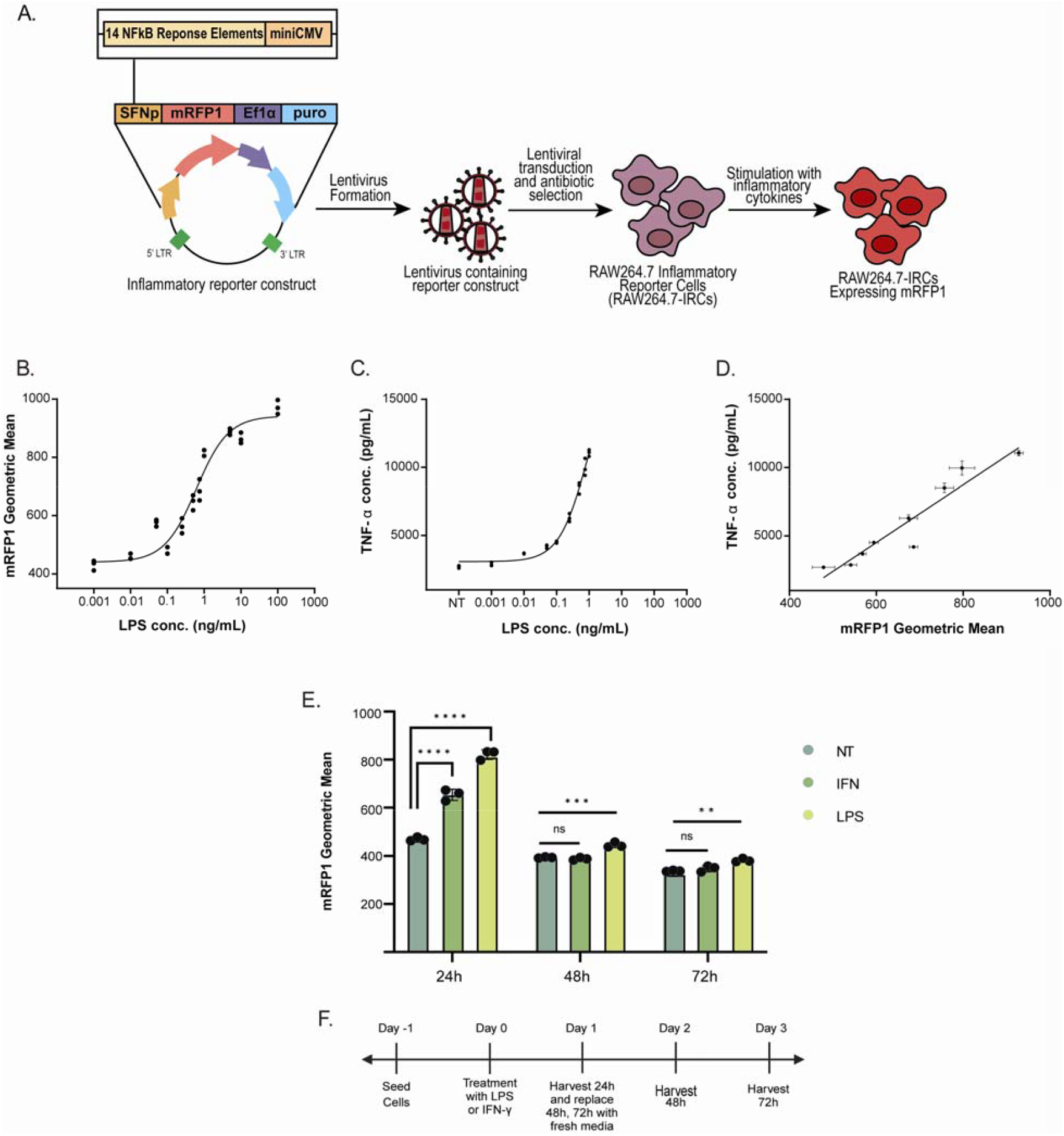
Characterization of RAW264.7-Inflammatory Reporter Cell line. **1A**. Overview of RAW264.7-IRC cell line creation. The inflammatory reporter construct, created using Gibson cloning, contained an mRFP1 fluorescent reporter under an inflammatory promoter, termed synthesis friendly inflammatory promoter (SFNp)^36^, and was encapsulated and transduced via lentivirus. The cells then underwent puromycin selection to create a homogenous population of cells. RAW264.7-IRCs were then stimulated using different cytokines and inflammatory agents to induce mRFP1 expression. **1B**. LPS dose curve showing range of mRFP1 fluorescence geometric mean values measured using flow cytometry across a range of LPS doses from 0.001 to 100 ng/mL. Individual data points are plotted for each biological replicate. Each dose was done in triplicate. Trend line calculated using the three-parameter dose-response curve, which yielded an EC50 = 0.61 ng/mL with an R^2^ value of 0.94. Dose-response curve line equation can be found in **Table S1**. **1C**. TNF-α concentration levels (pg/mL) in cell culture supernatant measured using ELISA. Cells were treated with LPS at concentrations ranging from 0.001 to 1 ng/mL for 24 hours. Each data point represents an average of 2 technical replicates. Trend line calculated using the three-parameter dose-response curve, which yielded an EC50 = 0.87 ng/mL with an R^2^ value of 0.987. Dose-response curve line equation can be found in Table S2. **1D**. Plot of mRFP1 fluorescent geometric mean values plotted against respective TNF-α concentration levels (pg/mL). *X*-values represent the average of the 3 biological replicates of TNF-α concentration (from Fig. 1C) and *y*-values represent the average of the 3 biological replicates of mRFP1 geometric mean (from Fig. 1B) ± SD. Trend line calculated using linear regression, *y* = 21.12*x*-8156, with an R^2^ value of 0.90. Linear regression parameters can be found in Table S3. **1E**. RAW264.7-IRC responsiveness. RAW264.7-IRCs were treated with LPS and IFN-γ for 24 hours, 24h cells were harvested for flow cytometry. Stimulation media was replaced with fresh media for the 48h and 72h samples. 24h post-removal of stimulation media, the 48h samples were harvested for flow cytometry, and the same was done for the 72h samples. Mean ± SD, with individual data points for each biological replicate. (^****^ p-value ≤ 0.0001, ^***^ p-value ≤ 0.001, ^**^ p-value ≤ 0.01, n.s. p-value > 0.05). **1F**. Timeline of treatment for RAW264.7-IRC responsiveness.

### Dynamic Range of Inflammatory Reporter

We assessed the dynamic range of the reporter in response to increasing levels of stimuli. The RAW264.7-IRCs were stimulated using LPS. RAW264.7-IRCs were seeded in 24-well plates. Dose levels ranged from 0.001 ng/mL to 100 ng/mL of LPS (**Fig. 1B**). 24 hours post-treatment, reporter cells were harvested in PBS, as described previously, and mRFP1 expression levels were quantified using flow cytometry. Cells were discarded after analysis. The data collected was analyzed using FlowJo™ v10.8 Software (BD Life Sciences). Cells were gated first to determine live cells by forward (FSC-A) and side scatter (SSC-A) and then FSC-A and FSC-H for single cells. Cells were then evaluated for mRFP1 expression (SSC v. mRFP1). Geometric mean values reported in the dose curve were taken from the single cell data. A dose response curve showing the dynamic range of the inflammatory reporter based on the geometric mean of the mRFP1 fluorescence seen in the single cell data set showed a dynamic response range from 0.1-1 ng/mL for the RAW264.7-IRCs (**Fig. 1B**). We also used ELISA to determine TNF-α levels in response to LPS activation for dose levels 0.001-1 ng/mL. Media from the RAW264.7-IRC cells used for the dose curve was collected before the cells were harvested and used to determine TNF-α levels released by the macrophages following 24 hours of LPS stimulation. TNF-α levels increased significantly over the dynamic range 0.1-1.0 ng/mL LPS, which aligns with the increasing inflammatory response seen by the inflammatory reporter across that range of LPS stimulation (**Fig. 1B&C**). To show the comparison between mRFP1 fluorescent geometric mean and TNF-α cytokine release by the RAW264.7-IRCs, the two values were plotted against each other and showed a linear correlation between the two values, with an R-squared value of 0.90 (**Fig. 1D**).

### Responsiveness of Inflammatory Reporter

The responsiveness of the inflammatory reporter was tested under LPS and interferon gamma (IFN-γ) stimulus. On day 0, RAW264.7-IRCs were treated with LPS and IFN-γ. 24 post-treatment, the 24h timepoint cells were harvested for flow cytometry, as described previously, and mRFP1 expression was quantified using the gating strategy described above. At this time, the media containing the stimuli for the 48h and 72h timepoint cells were removed and replaced with fresh media. 48h and 72h timepoint cells were harvested and mRFP1 expression quantified via flow cytometry (**Fig. 1E**). The inflammatory reporter showed a significant increase between the geometric mean intensity of the non-treated cells versus the treated cells (LPS, IFN-γ) 24 hours post-stimulus, as expected. At 48 and 72 hours post-treatment and the removal of the stimulus, there was a decrease in mRFP1 expression so that there was little to no significant difference seen between non-treated and previously treated RAW264.7-IRCs, showing that the reporter is responsive to the changing activation states of the macrophage (**Fig. 1F**).

### Phenotyping RAW264.7-IRCs using Immunostaining

Immunostaining and flow cytometry were performed to characterize the phenotypic surface expression 24 hours post-cytokine stimulation of the RAW264.7-IRCs. RAW264.7-IRC cells were treated with LPS at a concentration of 10 ng/mL for 24 hours prior to fixing and staining the cells for CD86 and CD206 to determine whether the cell populations exhibited a more inflammatory (M1) or anti-inflammatory (M2) phenotype based on cell surface markers CD86 is a common marker of classically activated mouse macrophages and CD206 is a common marker for alternatively activated mouse macrophages^38^. RAW264.7-IRCs stimulated with LPS showed a mixed phenotype displaying both the CD86 and CD206, like untreated RAW264.7-IRCs (**Fig. 2A**). Co-localization analysis of mean fluorescent intensity of mRFP1 with CD86 (blue) and CD206 (green) was performed using Fiji^39^. Co-localization analysis was performed for both non-stimulated RAW264.7-IRCs (**Fig. S1A**) and LPS-stimulated RAW264.7-IRCs (**Fig. S1B)**. For non-stimulated, mRFP1 fluorescence was minimized and there is more co-expression of both CD206 and CD86. For LPS-stimulated cells, there was an increase in CD86 mean fluorescent intensity, while CD206 decreased in mean fluorescent intensity. LPS stimulated cells exhibited an inflammatory phenotype, but mRFP1 expression is not as high as expected for this population although there is a trend towards increased mRFP1 expression. As the ratio of the fluorescent intensities of CD86:CD206 (a ratio of M1:M2) increased, mRFP1 fluorescence increased, but there was also a population with an increased inflammatory phenotype that do not have an increased mRFP1 fluorescence (**Fig. 2B**). Flow cytometry showcased an increase in both CD206 and CD86 after LPS stimulation but CD86 showed a larger population shift demonstrating its higher prevalence under inflammatory conditions (**Fig. 2C**).

**Figure 2.**
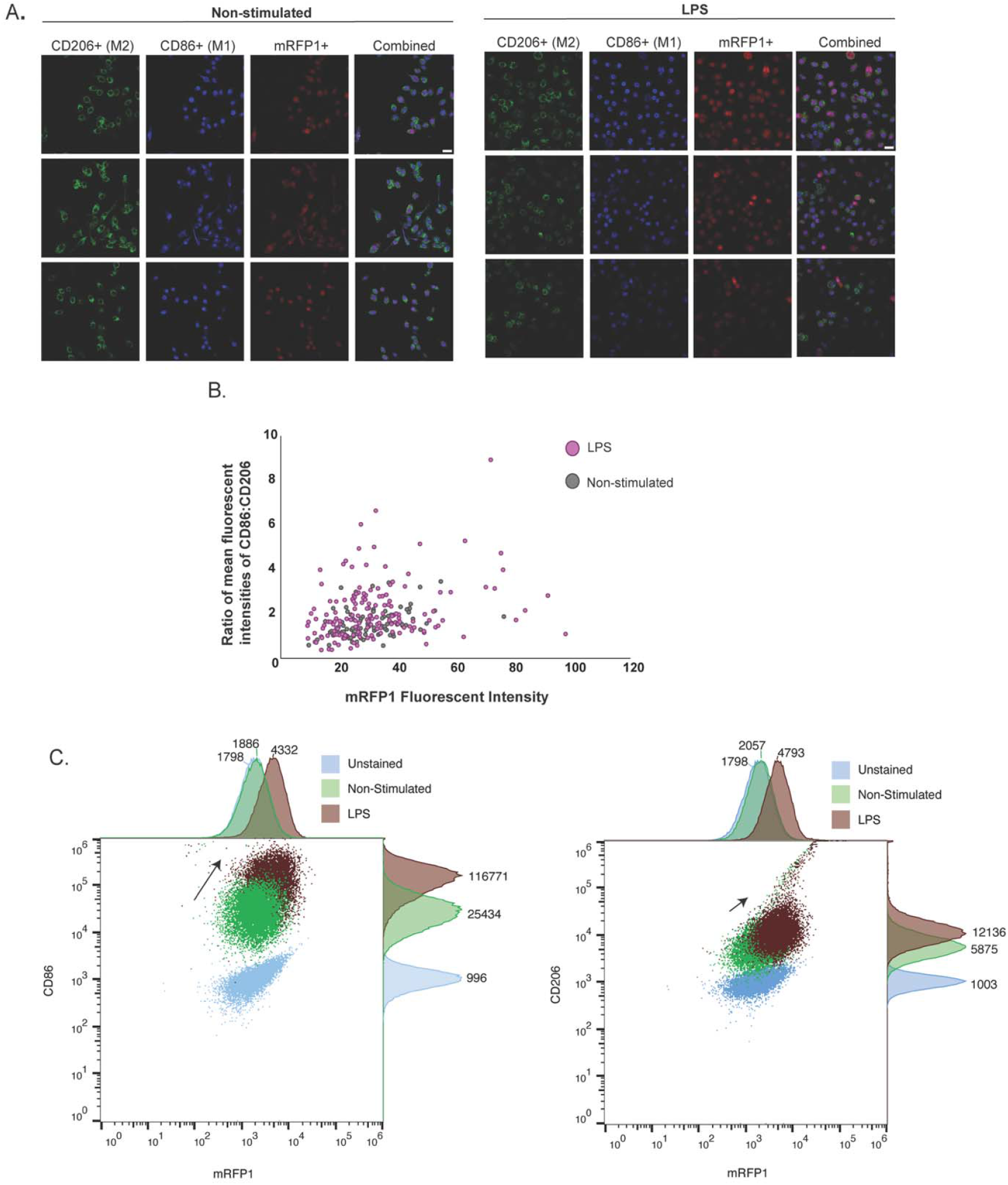
Immunostaining for Inflammatory versus Non-Inflammatory Phenotype in RAW264.7-IRCs. **2A**. Phenotyping via immunostaining of RAW264.7-IRCs for CD206 (green), a common anti-inflammatory marker, and CD86 (blue), a common inflammatory marker, co-localized with mRFP1 expression between no treatment and LPS treated cells (10ng/mL). Scale bars are 20 µm. **2B**. Ratio of mean fluorescent intensities of CD86:CD206 to mRFP1 fluorescent intensity as measured by confocal microscopy. **2C**. Phenotyping via flow cytometry of RAW264.7-IRCs for CD86 (left) and CD206 (right) alongside mRFP1 expression under three conditions: unstained, non-stimulated, and LPS treated (10 ng/mL). The black arrows showcase the magnitude of the cell population shift due to LPS stimulation.

### Inflammatory response of RAW264.7-IRCs to various viral and non-viral delivery methods

Viral and non-viral delivery vehicles were screened using RAW264.7-IRCs to determine the level of immune activation in response to the vehicles. For viral delivery, adeno-associated virus (AAV) serotypes 1, 2, 5, 8, and 9 at an MOI of 38,000, where MOI was determined based on the mass ratio of AAV to adenovirus, and adenovirus serotype 5/35 (Ad5/35) at an MOI of 1,000 were used to transduce the inflammatory reporter cells with a CMV promoter-driven green fluorescent protein (GFP). For non-viral delivery, Lipofectamine^TM^ 2000 and TransIT-X2® Dynamic Delivery System (Mirus bio), a polymer with proprietary components that aid in cell uptake, were used to deliver a plasmid encoding CMV promoter-driven GFP to RAW264.7-IRCs. Cells receiving viral delivery were analyzed using flow cytometry to quantify GFP and mRFP1 levels at both 24 and 48 hours post-transduction. Cells were harvested for flow cytometry and mRFP1 and GFP fluorescence was quantified. Cells transfected with non-viral delivery vehicles were harvested and GFP and mRFP1 fluorescence analyzed via flow cytometry at 48 hours post-transfection, per the respective manufacturer’s protocol. Using both GFP and mRFP1, the dual-reporter assay elucidated both delivery efficiency based on GFP levels and inflammatory response based on mRFP1 levels. Following flow cytometry, data was analyzed using FlowJo™ Software with the same gating process mentioned above with an additional step of gating on a quadrant of GFP vs. mRFP1. **Figure 3A** shows the percentage of cells in each quadrant (q1: mRFP1-/GFP-; q2: mRFP1+/GFP-; q3: mRFP1+/GFP+; q4: mRFP1-/GFP+). There was no observable difference between AAV2, AAV5, AAV8, AAV9, and the control in inflammatory response. AAV2, AAV5, AAV8, and AAV9 also showed no GFP delivery to the RAW264.7-IRCs. AAV1 showed a slight increase in GFP delivery with an average of 25.9% GFP+ cells. As expected, Ad5/35 showed a significant increase in the efficiency of GFP delivery with an average of 56.4% GFP+ cells, but interestingly, the delivery vehicle was only slightly more inflammatory than others with an average 75.4% mRFP1+ cells compared to 61.2% for reporter control. Similarly, to what was seen in **Figure 1E**, AAV2, AAV5, AAV8, and AAV9 showed a relative decrease in mRFP1+ cells, illustrating a return to an alternatively activated state. Ad5/35 had the most significant GFP expression with an average of 56.4% and 45.1% GFP+ cells at 24h and 48h, respectively, and increased mRFP1 expression compared to reporter control at 24h (75.4% vs. 61.2%). Interestingly, the reporter control maintained its inflammatory response compared to treatment groups which averaged a 22.5% decrease in %mRFP1+ from 24h to 48h. TransIT-X2 delivery of GFP resulted in efficient GFP delivery (35.8% GFP+), but also significantly increased inflammatory response (95.2% mRFP1+) versus reporter control at 48h (67.9% mRFP1+). Lipofectamine 2000 delivery resulted in an inflammatory response like TransIT-X2 delivery, with 96.2% mRFP1+ and less efficient GFP delivery with 20.5% GFP+. Delivery data was plotted to show the ratio of efficient delivery to inflammatory response using the fluorescence geometric mean values for both GFP and mRFP1 (GFP Geometric Mean:mRFP1 Geometric Mean) (**Fig. 3B**). Geometric means were used to represent the data due to the wide range of fluorescent intensities in the sample cells and to lessen the effects of extreme outliers in the flow data, namely with the non-viral delivery, such as with TransIT-X2 (**Fig. 3C**). Again, Ad5/35 presents the only viable option for both efficient delivery and a lower inflammatory response, whereas with non-viral delivery, Lipofectamine 2000 and TransIT-X2, there is a higher delivery efficiency but also significantly increased inflammatory response to these modes of delivery. Interestingly, when graphing the geometric means, the reporter control shifts drastically from the 24h to the 48h data set and shows an increased GFP geometric mean of 198.8 vs. 380.8, respectively, and increased mRFP1 geometric mean of 380.8 vs. 690.4, indicating a heightened inflammatory state. To better understand the source of variability, we sought to measure the effects of confluency on mRFP1 expression and inflammatory state of the RAW264.7-IRCs (**Fig. S2-3**). To determine the effects of confluency on the baseline mRFP1 fluorescence, RAW264.7-IRCs were seeded in 6-well plates to achieve a range of cell densities. Cells were then harvested at 24h and 48h post-seeding for flow sorting. The results showed a remarkably consistent trend at 24 and 48h for a gradual decrease in %mRFP1+ and mRFP1 fluorescent geometric mean for low cell seeding densities followed by a dramatic increase in %mRFP1+ and mRFP1 fluorescent geometric mean. Overall, depending on the time points and confluency, the range for %mRFP1 positive was around 20% to 40% and geometric mean ranged from 200 to almost 400 across both time points and different cell densities (**Fig. S2**).

**Figure 3.**
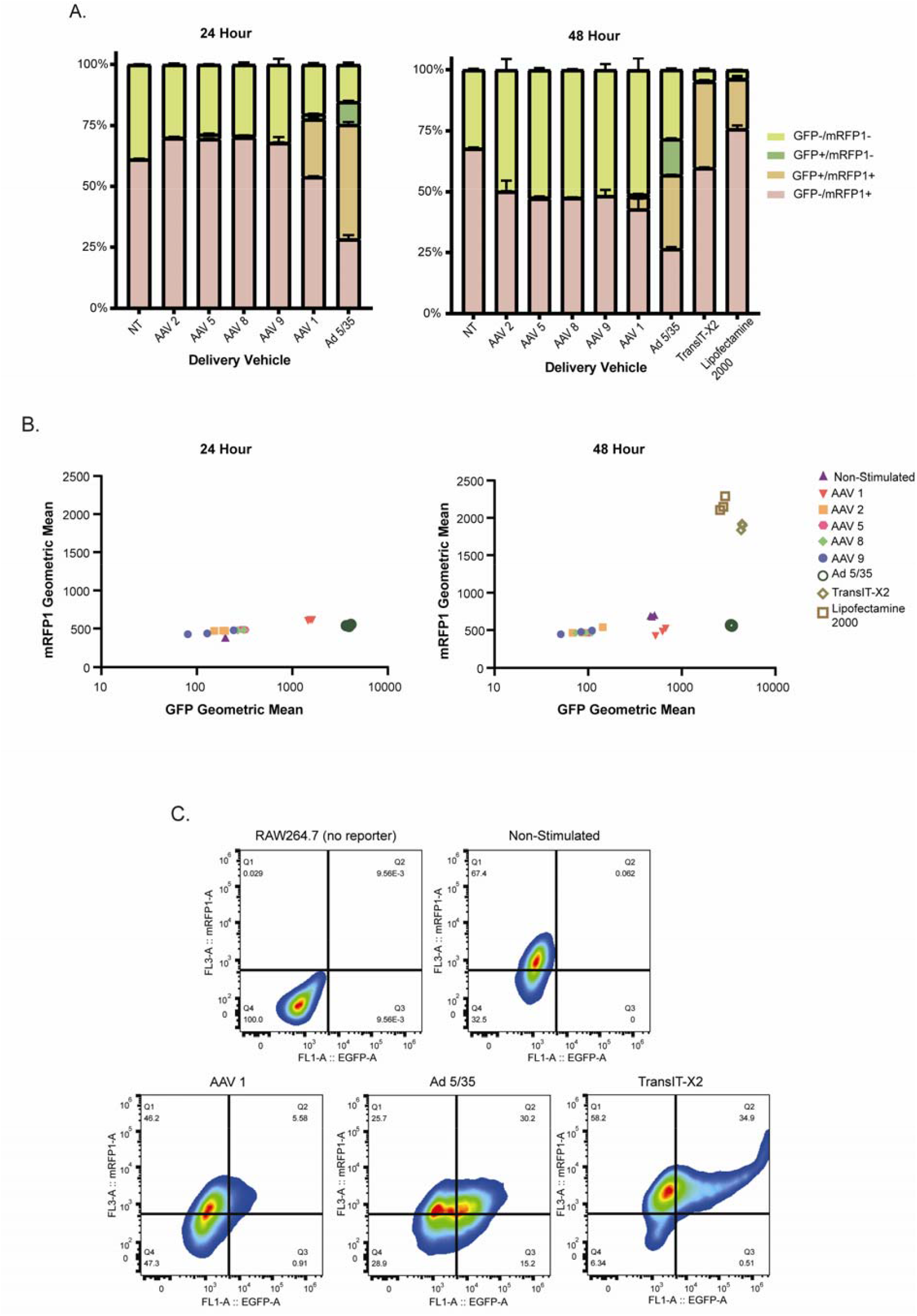
Evaluation of Inflammatory Response and Delivery Efficiency of Viral and Non-Viral Delivery Vehicles using RAW264.7-IRCs. **3A**. Evaluation of viral and non-viral delivery methods using a dual-reporter assay at 24 and 48 hours post-delivery. Efficiency of delivery and inflammatory response were determined by using different viral (AAV 1, 2, 5, 8, 9 and Ad5/35) and non-viral (TransIT-X2 and Lipofectamine 2000) delivery methods carrying a GFP payload to RAW264.7-IRCs at timepoints of 24 hours and 48 hours post-delivery. %GFP+ representing efficient delivery and %mRFP1+ representing inflammatory response. Percentages represent the average of 3 biological replicates. Each bar represents the average percentages from 3 biological replicates. TransIT-X2 and Lipofectamine 2000 were only evaluated at 48 hours post-transfection per manufacturer protocol. Raw data points can be found in Table S4. **3B**. Evaluation of the geometric mean of GFP to mRFP1 at 24 and 48 hours post-delivery. The data represented here is the same samples in 2A. Individual data points were plotted for each biological replicate. Y-axes between 24 and 48 hours is different. Raw data points can be found in Table S4. Statistics done using One-Way Anova and Tukey’s Post-Hoc Analysis can be found in Table S5 for 24 hour and S6 for 48 hour. **3C**. Representative cell populations from dual-fluorescence reporter assay. Populations represented here are RAW264.7 cells with no reporter to set the negative control for GFP/mRFP1, non-treated RAW264.7-IRCS (NT) to determine basal mRFP1 expression among reporter cells, and then three representative populations from RAW264.7-IRCs treated with AAV 1, Ad 5/35, and TransIT-X2, respectively. Gating strategy for dual-fluorescence assay can be found in Figure S2.

## Discussion

As therapeutics continue to advance, it is important to continually develop tools that aid in their rapid development. Macrophages represent a compelling target for regenerative and immunomodulatory therapeutics due to their innate nature as they both defend the body and facilitate the return to and maintenance of homeostasis^9^. It is essential to know how macrophages will respond to different therapeutics including small molecule drugs, gene delivery vehicles, or new vaccines. In the early development stage, high-throughput screening techniques, such as the one described here, could provide simultaneous results on gene delivery and inflammatory responses for candidate vectors.

The reporter developed here can be used to investigate the role of inflammation through the NF-κB activation pathway, which is important in regulating the expression of various inflammatory genes, including TNF-α^40–42^. The RAW264.7-IRCs utilize the NF-κB activation pathway through the 14 NF-κB response elements to promote mRFP1 expression. This in turn relays the relative amount of inflammation due to activated NF-κB, that has translocated to the nucleus. We were able to confirm the inflammatory status of the RAW264.7-IRCs and their relative mRFP1 expression to TNF-α released by the cells at different LPS doses (**Fig. 1B&C**). The inflammatory reporter is both responsive and titratable. The LPS dose curve and TNF-α concentration curve corresponding to the same LPS doses, ranging from 0.001 to 1 ng/mL, show that the mRFP1 expression is strongly correlated with the common inflammatory marker, TNF-α. The reporter shows a responsiveness to its environment and can recover from inflammatory stimulus back to baseline levels, which is in line with the plastic nature of macrophages (**Fig. 1E**). Interestingly, through immunostaining for macrophages markers CD86 (M1) and CD206 (M2), we saw similar phenotypes with both markers expressed on both non-treated and LPS treated RAW264.7-IRCs, which could be due to the nature of the RAW264.7 cell line to exist as phenotypically diverse at baseline. RAW264.7 cells are an immortalized cell line and experience a more homogenous NF-κB response than primary macrophages. RAW264.7 cells exhibit a lower basal activated (nuclear) NF-κB level at about 5-10% of total cellular NF-κB compared to bone-marrow derived mouse macrophages (BMMs) with a higher basal activated NF-κB level at 25-35% of total cellular NF-κB. Also of note, RAW264.7s only reached 50-60% of total cellular NF-κB translocation to the nucleus upon LPS stimulation whereas BMMs achieved activated NF-κB levels of 80-90% of total cellular NF-κB^43^. The basal levels of activated NF-κB could also contribute to the levels of mRFP1 expression observed in non-treated RAW264.7-IRCs, especially in those seeded for an optimal confluency.

In order to draw meaningful conclusions, a non-stimulated RAW264.7-IRC control should be used for every experiment to normalize the inflammation experienced by the cells due to environmental factors such as confluency, which was determined to be a contributor to mRFP1 expression and can result in a range of basal mRFP1 expression of 20-40% mRFP1+ cells or a geometric mean for mRFP1 fluorescence ranging from 250-400 (**Fig. S2-S3**). The lower geometric mean values seen at lower cell confluency could be due to its entrance into the cell cycle which has been shown to suppress inflammation^44^. Additionally, the difference between 24 and 48 hrs. may be due to the increase in cell density after 1-2 doubling times for RAW 264.7 cells or the length of time the macrophages are being signaled to undergo mitosis. These macrophages represent a very heterogenous population and tend to change size and morphology upon stimulation. Due to this, the geometric mean was used to account for extremes in the different populations and compare the bulk of the phenotypes seen in each treatment group during flow cytometry analysis. The geometric mean weights the mean fluorescent intensity based on the bulk cell population and gives less weight to the upper extremity of the distribution. This was especially important when comparing treatments such as TransIT-X2 and Lipofectamine 2000 to the other groups as a small portion of RAW264.7-IRCs expressed much higher levels of mRFP1 (**Fig. 3C**).

After characterizing the inflammatory reporter cell line, the RAW264.7-IRCs were used for a dual-reporter assay to determine inflammatory response in macrophages to different viral and non-viral delivery vehicles and delivery efficiency. The various delivery vehicles had an array of effects on activation of the mouse macrophages, ranging from some viral vectors showing zero effect on the macrophages, with basal mRFP1 levels and no GFP expression, to the non-viral vectors showing both increased delivery and greatly increased inflammation. Using the RAW264.7-IRCs to investigate the effects of various delivery vectors, both viral and non-viral, we saw that AAV serotypes 1,2, 5, 8, and 9 had no effect on either delivery or inflammatory response by the macrophages. AAV1 showed an increase in both GFP and mRFP1 fluorescence compared to the other AAV serotypes tested. In comparing the transduction efficiency of AAV1 across timepoints and the drastic change in the control RAW264.7-IRCs from 24 to 48 hours, it is difficult to draw conclusions on the effectiveness of that specific vector. As expected, Ad5/35 had the best GFP delivery for the macrophages due to the engineering of the adenovirus to target CD46, which is expressed by macrophages, and increasing the level of uptake in those cells^16,45^. Ad5/35 is used in the production of chimeric antigen receptor macrophages (CAR-Ms) which are macrophages that are modified *ex vivo* before being transduced back into the patient to target cancers, and as such must maintain an inflammatory (M1) macrophage phenotype to induce an immune response in the tumor microenvironment. The issue with the viral vectors that lead to more efficient delivery is the inflammatory response associated with their use. For some applications, such as with CAR-Ms for immunotherapy in cancer, an inflammatory phenotype is desirable due to the immunosuppressive nature of tumors and a need to mount a response to the tumor^46–48^. Although this may be the goal for some therapeutics, other approaches may require an anti-inflammatory phenotype for the macrophages post-delivery with viral or non-viral vehicles. This approach is especially important for a regenerative medicine setting where inflammation, both acute and chronic, is usually the cause for prolonged healing processes or pathologic fibrosis^3,49,50^. Either way, the reporter developed here can provide insight into inflammatory or anti-inflammatory nature of delivery vehicles and aide in high-throughput screening for rapid candidate development.

## Conclusions

In summary, this inflammatory reporter provides a sensitive and dynamic screening tool to determine the response to different stimuli and reveal information on inflammatory responses by macrophages. As for future studies, this inflammatory reporter can be used in different applications when looking at inflammatory response or processes related to NF-κB activation. One study used an NF-κB reporter for CRISPR knockout screening to determine different genes important to inflammatory response and essential for viability in macrophages^35^. This reporter could also provide insights for small molecules screening on inflammation^51^ or elucidate the effects of vaccines on macrophages during development^52,53^. Overall, this work demonstrates a well-characterized screening technique for inflammatory response in macrophages that can be useful in gene delivery and adoptive cell therapy.

## List of Abbreviations

AAV: adeno associated virus
AdV: adenovirus
GFP: green fluorescent protein
RAW264.7-IRC: RAW264.7 inflammation reporter cells
NF-κB: nuclear factor-kappa B
TNF-α: tumor necrosis factor alpha
CD86: cluster of differentiation 86
CD206: cluster of differentiation 206
CAR-Ms: chimeric antigen receptor macrophages
TAMs: tumor associated macrophages
LPS: lipopolysaccharide
SFNp: synthesis-friendly inflammation-inducible promoter

## Declarations

### Ethics approval and consent to participate

n/a

### Consent for publication

n/a

### Availability of data and materials

The datasets generated and/or analyzed during the current study are available in the Github repository, https://github.com/chrisnelsonlab/DualReporterMacrophage1. Plasmids have been deposited with the non-profit repository addgene.

### Author contributions

AI and CN designed experiments and wrote the manuscript. AI performed experiments, analyzed data, and created figures. SA and AS performed flow cytometry experiments and analysis. SB performed immunostaining and confocal imaging. TJM assisted with confocal analysis, experimental design for macrophage imaging, and manuscript revisions. VK aided in confocal imaging.

## Acknowledgments

We acknowledge members of the Nelson and Muldoon labs for helpful conversations on macrophage biology and general laboratory support.

