## Supplemental Information for "A Dual-Fluorescence Assay for Gene Delivery Vehicle Screening in Macrophages with an Inflammation-Inducible Reporter Construct"

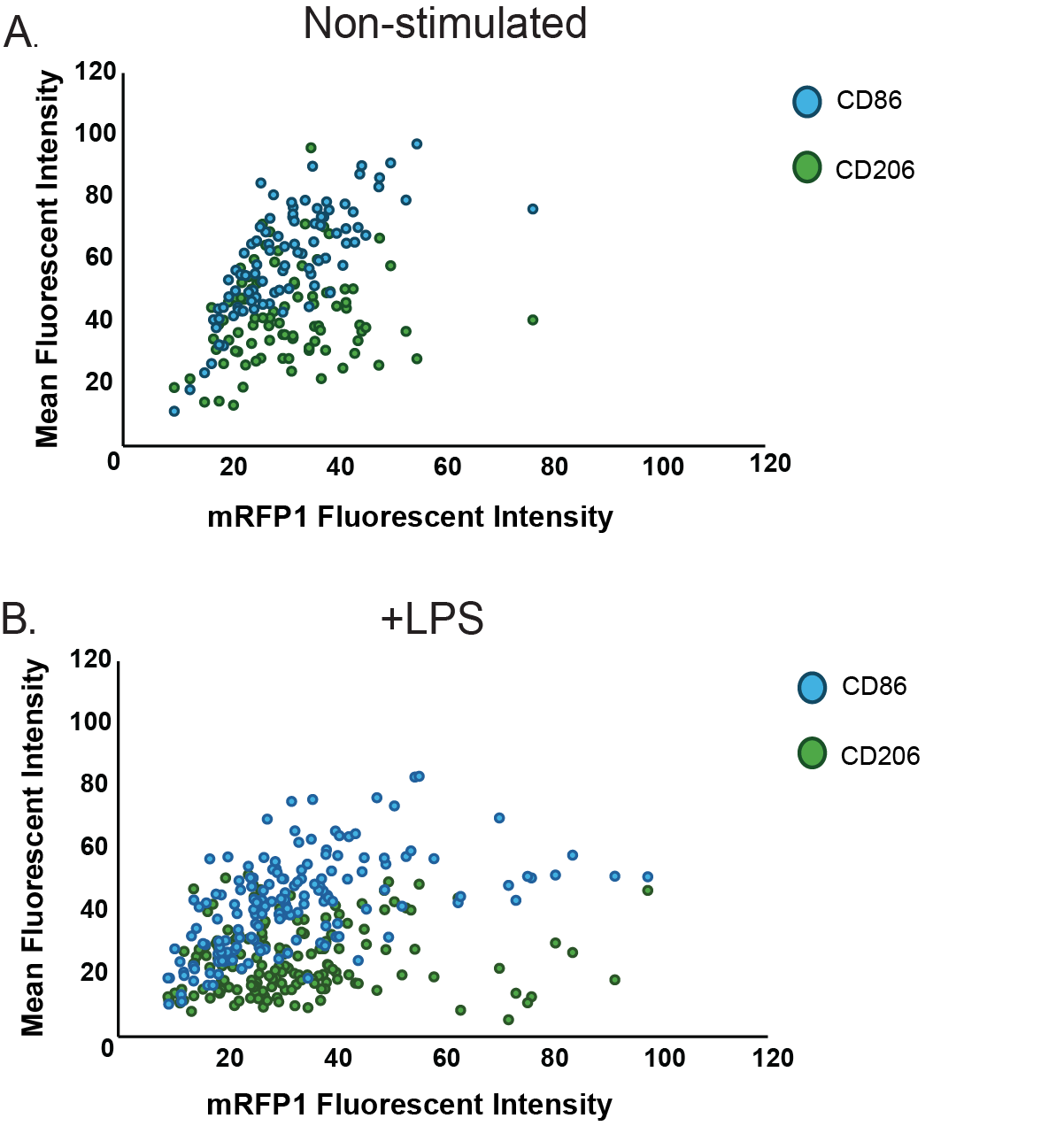

**Figure S1 |** Confocal colocalization measurement for mRFP1 fluorescent intensity and CD86 or CD206 intensity for A) non-stimulated IRCs and B) LPS stimulated IRCs.

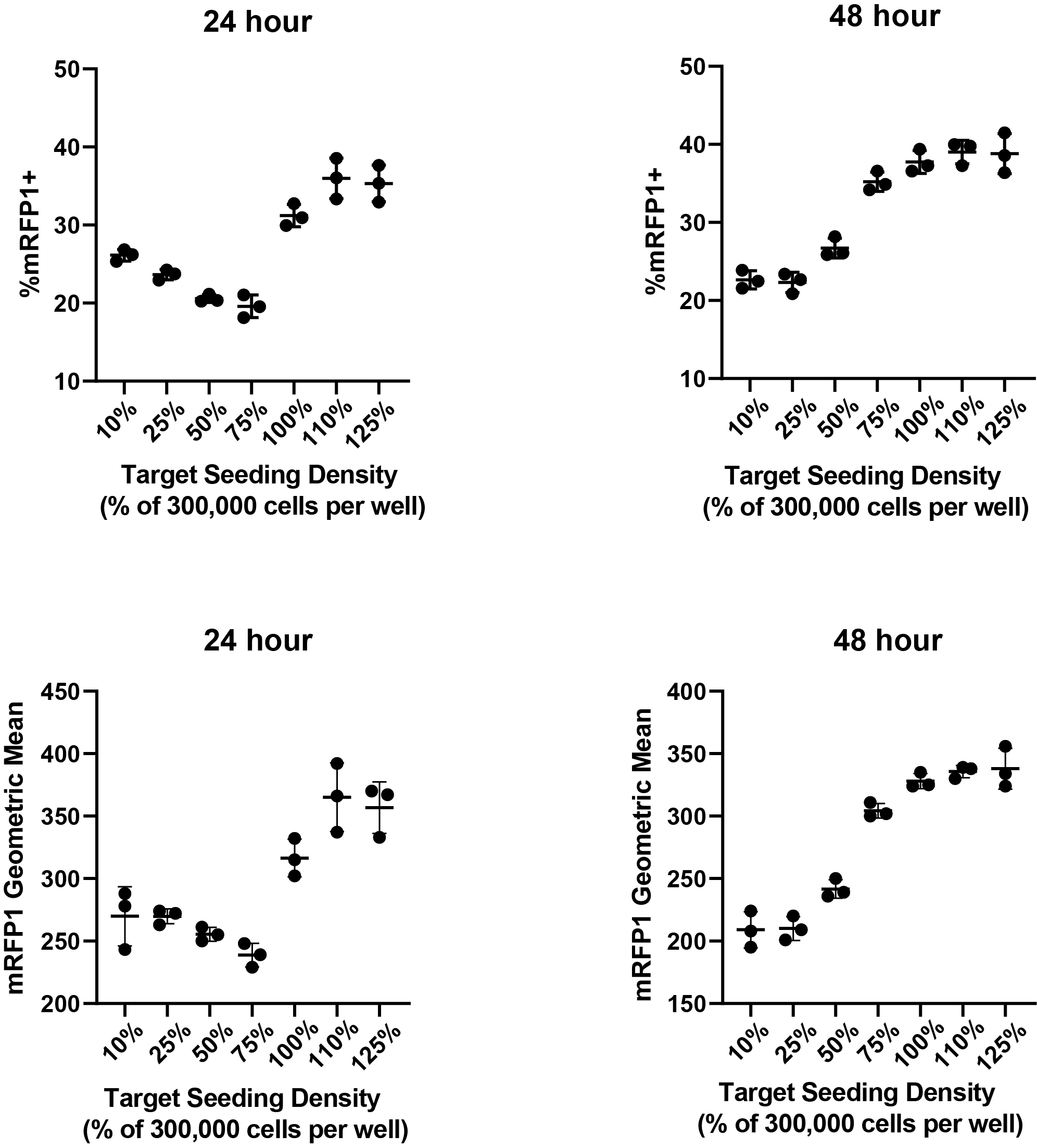

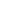

**Figure S2 | Effects of confluency and time in culture on mRFP1 expression.** Cells were seeded in 6-well plates at a percentage of the seeding density suggested for 6-well plates (300,000 cells/well is considered 100%). Cells were allowed to expand for 24 and 48-hours respectively, with cells in the 48-hour time point seeded at half the seeding density of the 24-hour cells. Cells were imaged to show confluency and then harvested for flow cytometry. mRFP1 fluorescence presented here as both %mRFP1+ and geometric mean to show the range of mRFP1 expression that is observed through the no treatment groups. Individual data points represent a single biological replicate, experiments were run in triplicate.

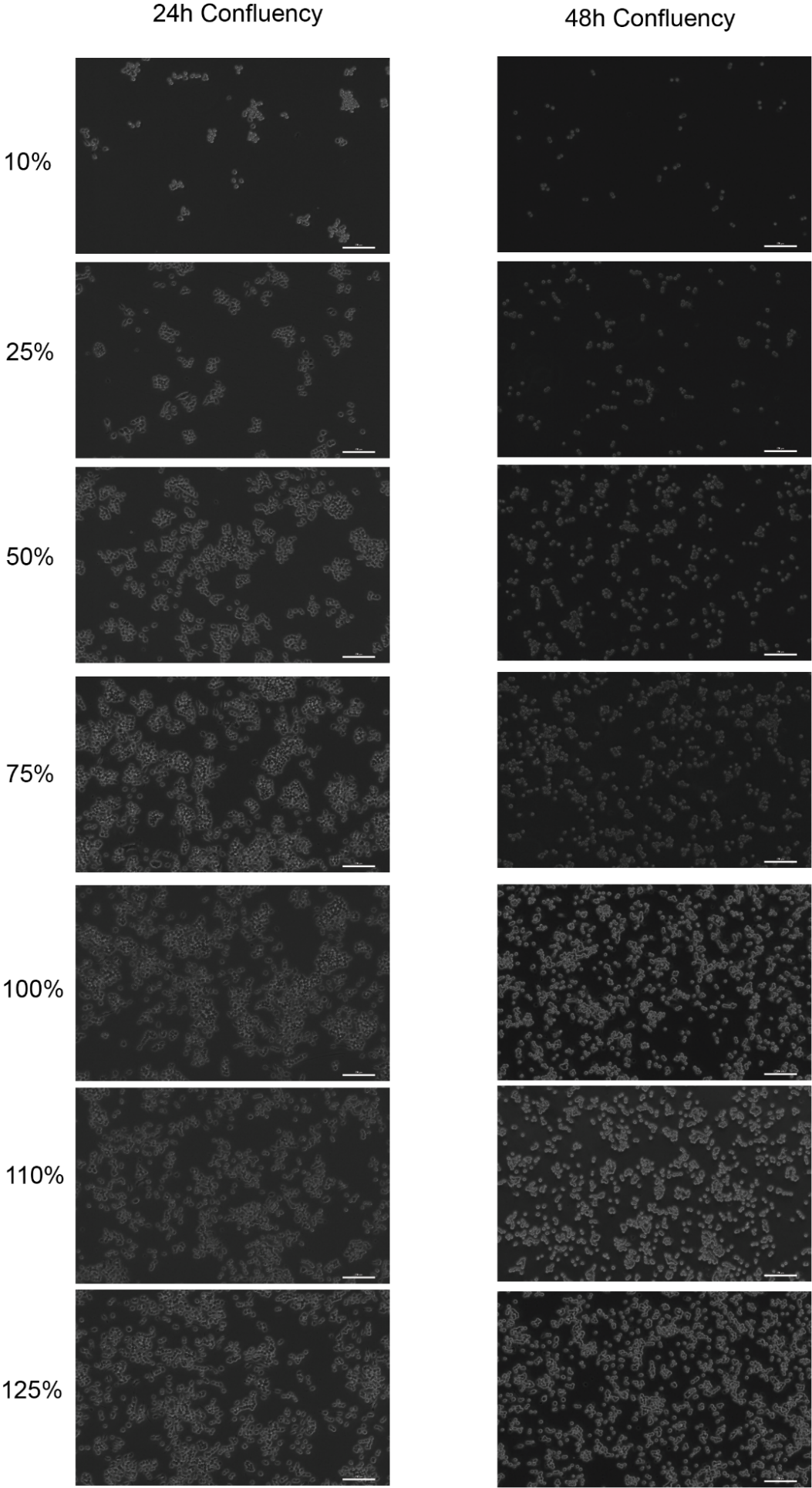

**Figure S3 | Representative images for confluency study -** For corresponding images of cell confluency, mages were taken using the 10X objective. Scale bars are 250 µm.

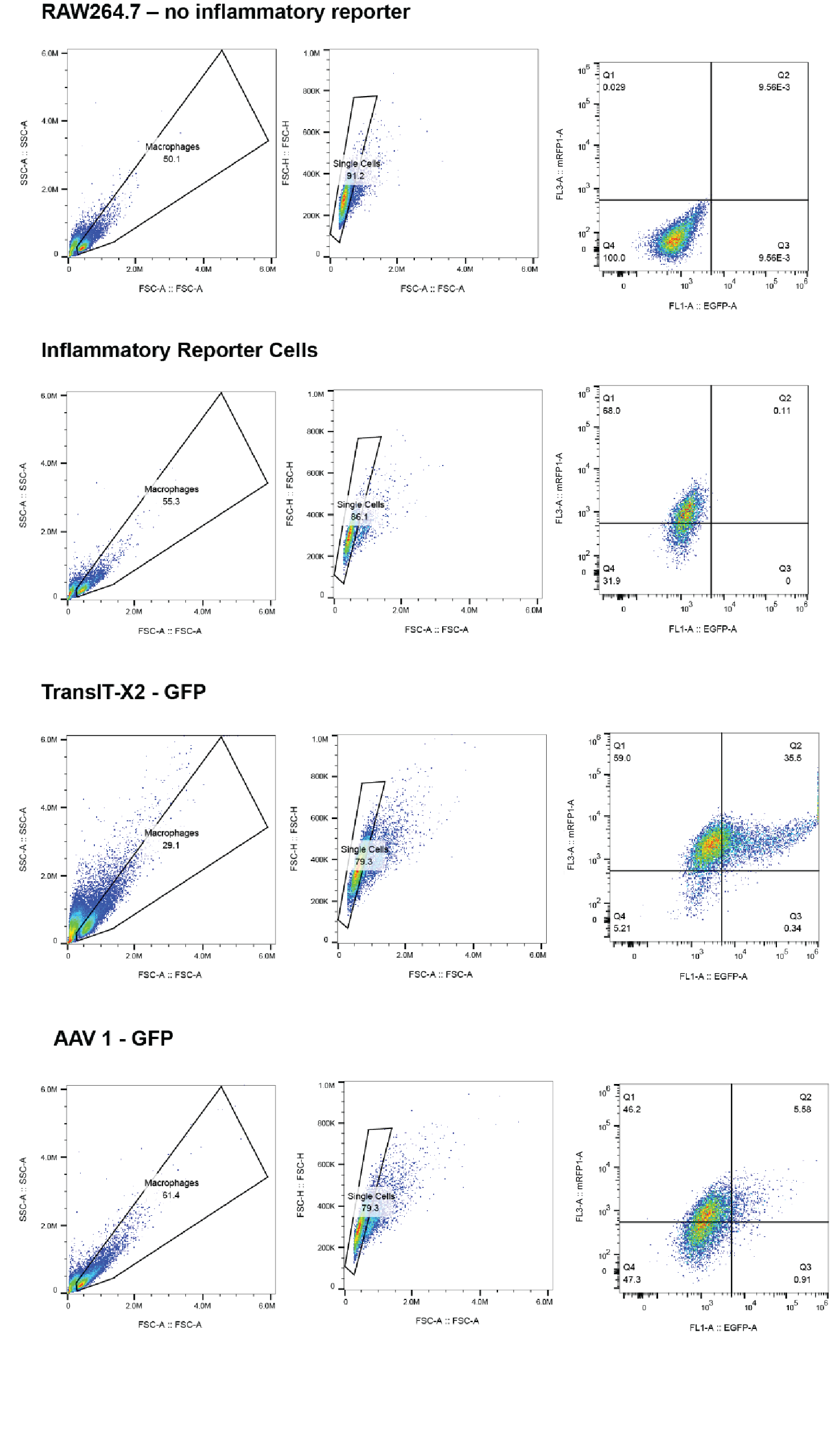

**Figure S4.** **Gating strategy for dual-reporter assay (GFP and mRFP1).** Cells were harvested and resuspended in PBS for flow cytometry. RAW264.7 cells without the mRFP1 inflammatory reporter were always used to set the initial gate for %mRFP1-. The gates were then checked against the treated cells to make sure all cells were included as there would be a drastic change in the range of cell size following treatment. Cells were gated FSC-A vs SSC-A to capture live macrophages from cell debris and then gated on FSC-A vs FSC-H for single cells. Cells were then gated on a quadrant of EGFP vs mRFP1. Using the RAW264.7 cells with no reporter, the quadrant gate was set so that population represented the total %mRFP1-, %GFP-. Gating was done after flow cytometry data was gathered using the FlowJo™ Software.

**Table S1**. LPS Dose Curve – Curve Fitting with Agonist vs. Response (Fig. 1B)

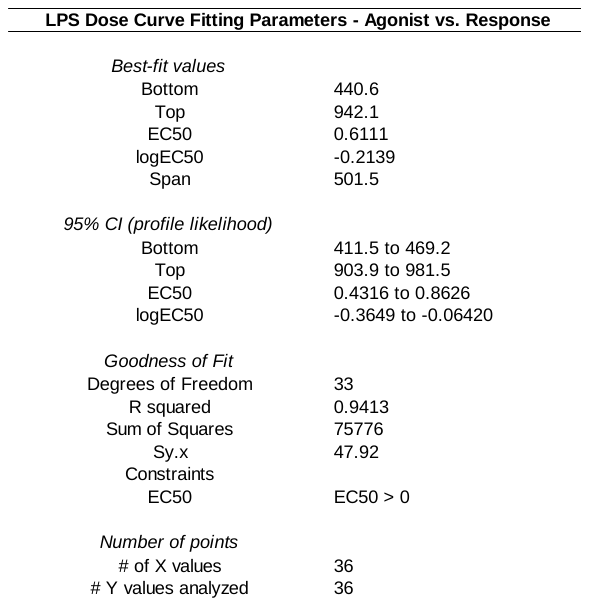

**Table S2**. TNF-a Concentration Curve Fitting Parameter – Agonist vs Response (Fig. 1C)

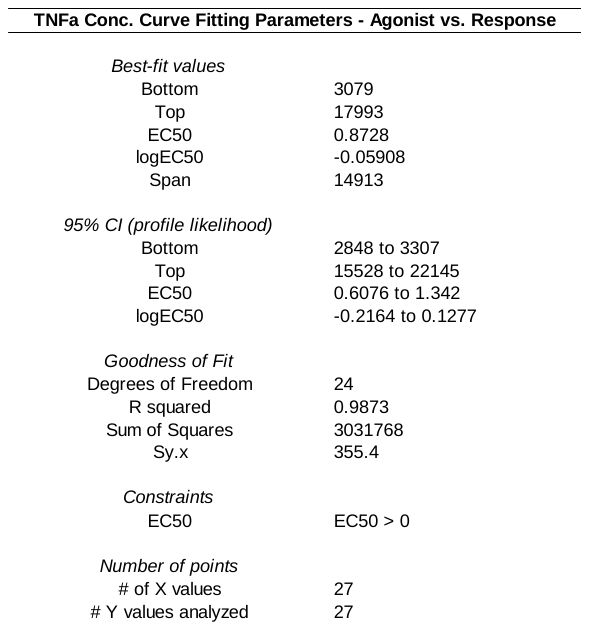

**Table S3**. TNF-α Concentration vs. mRFP1 Expression – Linear Regression Parameters (Fig. 1D)

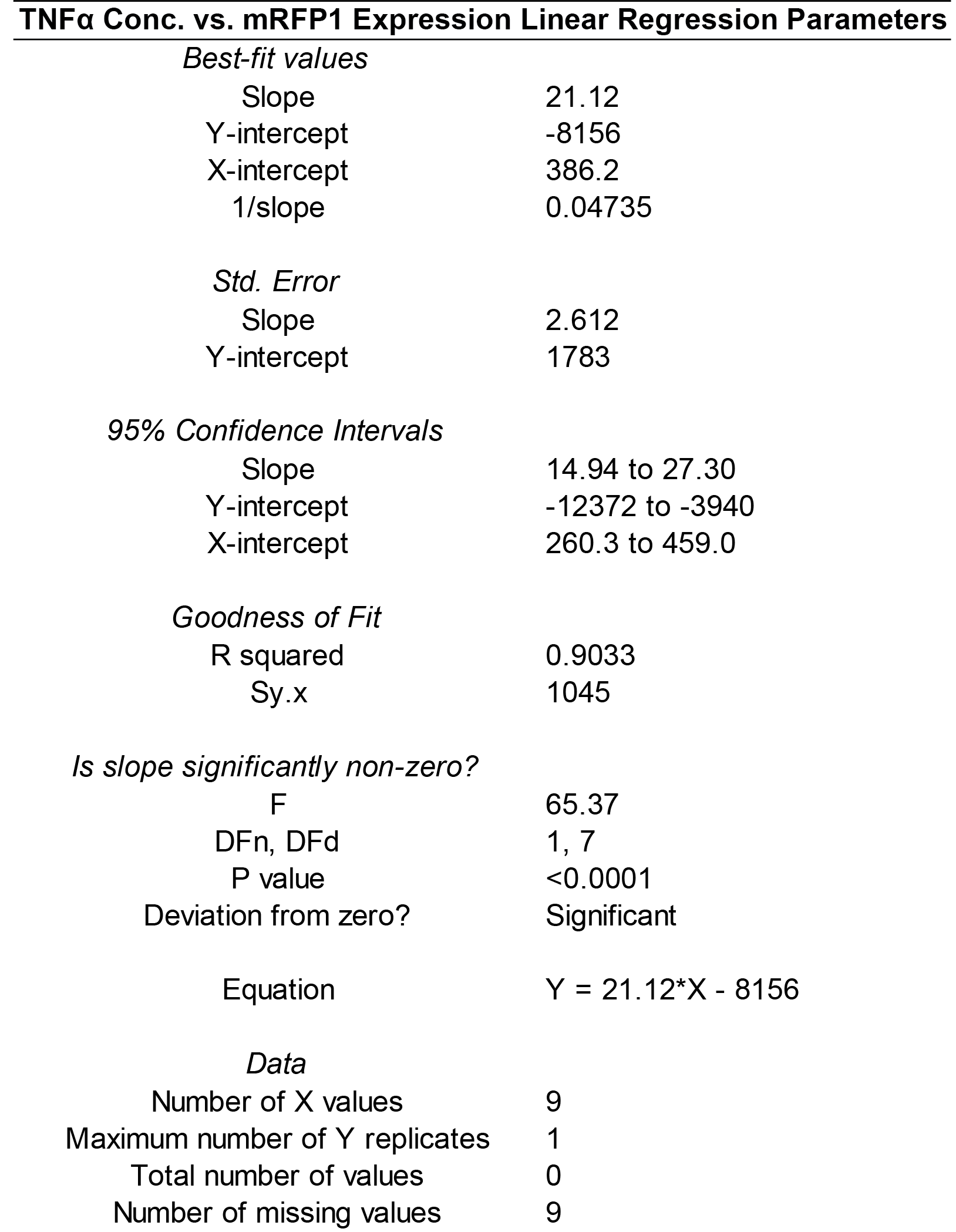

**Table S4.** Raw data for Fluorescence Geometric Mean and %GFP/mRFP1 Quadrant Populations from Flow Cytometry (Figures 2 A&B)

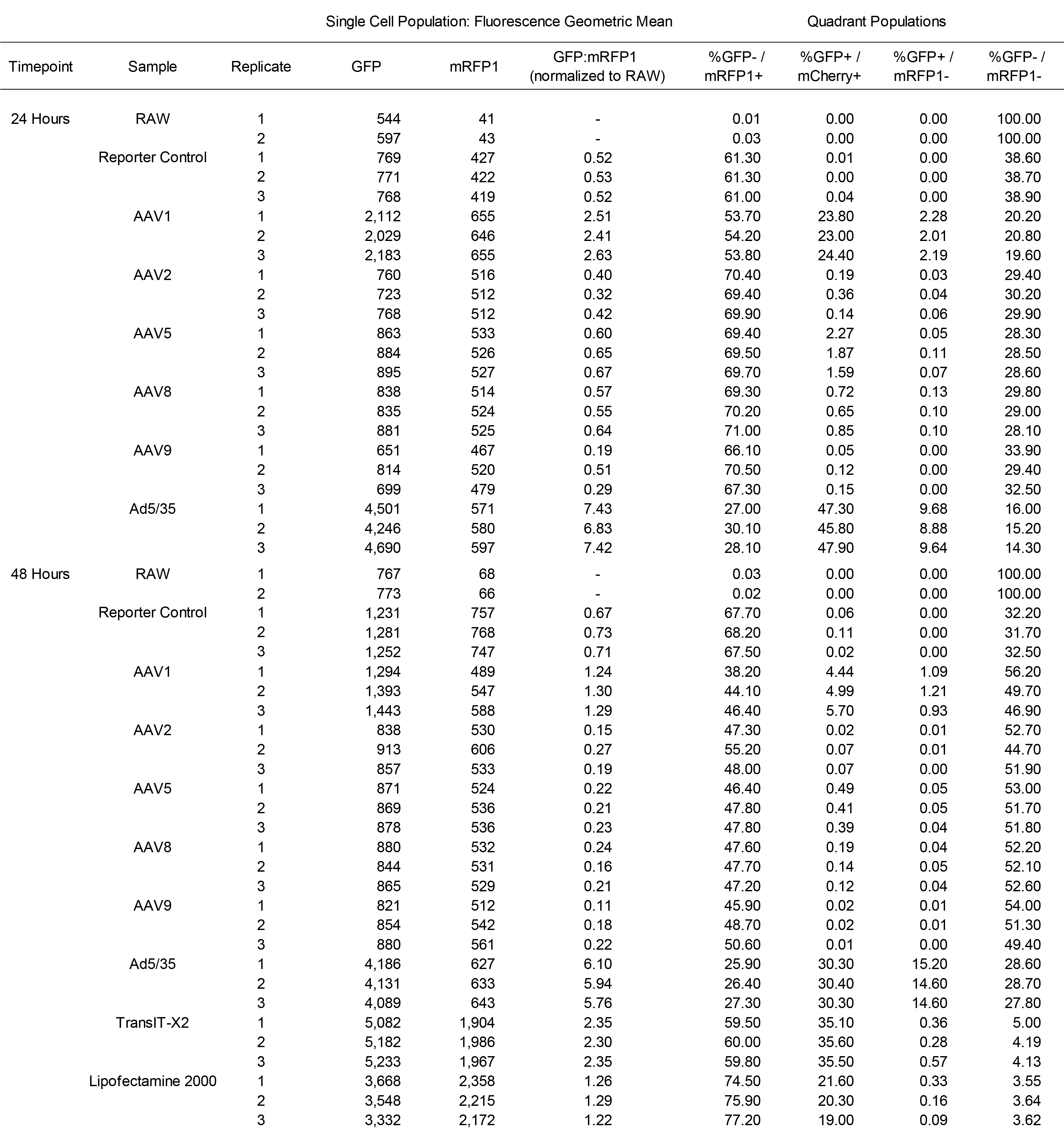

**Table S5.** Summary statistics using One-Way Anova and Tukey’s Post-Hoc Analysis for 24-hour delivery vehicle GFP:mRFP1 ratio, GFP and mRFP1 fluorescent geometric means normalized to RAW264.7 cells with no GFP/mRFP1 (Figures 2 B)

| **Sample** | **Comparator** |  | **P-value Summary** |
| --- | --- | --- | --- |
| AAV1 | Reporter Control |  | **** |
|  | AAV2 |  | **** |
|  | AAV5 |  | **** |
|  | AAV8 |  | **** |
|  | AAV9 |  | **** |
|  | Ad5/35 |  | **** |
| AAV2 | Reporter Control |  | ns |
|  | AAV1 |  | **** |
|  | AAV5 |  | ns |
|  | AAV8 |  | ns |
|  | AAV9 |  | ns |
|  | Ad5/35 |  | **** |
| AAV5 | Reporter Control |  | ns |
|  | AAV1 |  | **** |
|  | AAV2 |  | ns |
|  | AAV8 |  | ns |
|  | AAV9 |  | ns |
|  | Ad5/35 |  | **** |
| AAV8 | Reporter Control |  | ns |
|  | AAV1 |  | **** |
|  | AAV2 |  | ns |
|  | AAV5 |  | ns |
|  | AAV9 |  | ns |
|  | Ad5/35 |  | **** |
| AAV9 | Reporter Control |  | ns |
|  | AAV1 |  | **** |
|  | AAV2 |  | ns |
|  | AAV5 |  | ns |
|  | AAV8 |  | ns |
|  | Ad5/35 |  | **** |
| Ad5/35 | Reporter Control |  | **** |
|  | AAV1 |  | **** |
|  | AAV2 |  | **** |
|  | AAV5 |  | **** |
|  | AAV8 |  | **** |
|  | AAV9 |  | **** |

**Table S6.** Summary statistics using One-Way Anova and Tukey’s Post-Hoc Analysis for 48 hour timepoint delivery vehicle GFP:mRFP1 ratio with GFP and mRFP1 fluorescent geometric means normalized to RAW264.7 cells with no GFP or mRFP1 (Figures 2B)

| **Sample** | **Comparator** |  | **P-value Summary** |
| --- | --- | --- | --- |
| AAV1 | Reporter Control |  | **** |
|  | AAV2 |  | **** |
|  | AAV5 |  | **** |
|  | AAV8 |  | **** |
|  | AAV9 |  | **** |
|  | Ad5/35 |  | **** |
|  | TransIT-X2 |  | **** |
|  | Lipofectamine 2000 |  | ns |
| AAV2 | Reporter Control |  | **** |
|  | AAV1 |  | **** |
|  | AAV5 |  | ns |
|  | AAV8 |  | ns |
|  | AAV9 |  | ns |
|  | Ad5/35 |  | **** |
|  | TransIT-X2 |  | **** |
|  | Lipofectamine 2000 |  | **** |
| AAV5 | Reporter Control |  | **** |
|  | AAV1 |  | **** |
|  | AAV2 |  | ns |
|  | AAV8 |  | ns |
|  | AAV9 |  | ns |
|  | Ad5/35 |  | **** |
|  | TransIT-X2 |  | **** |
|  | Lipofectamine 2000 |  | **** |
| AAV8 | Reporter Control |  | **** |
|  | AAV1 |  | **** |
|  | AAV2 |  | ns |
|  | AAV5 |  | ns |
|  | AAV9 |  | ns |
|  | Ad5/35 |  | **** |
|  | TransIT-X2 |  | **** |
|  | Lipofectamine 2000 |  | **** |

| **Sample** | **Comparator** |  | **P-value Summary** |
| --- | --- | --- | --- |
| AAV9 | Reporter Control |  | **** |
|  | AAV1 |  | **** |
|  | AAV2 |  | ns |
|  | AAV5 |  | ns |
|  | AAV8 |  | ns |
|  | Ad5/35 |  | **** |
|  | TransIT-X2 |  | **** |
|  | Lipofectamine 2000 |  | **** |
| Ad5/35 | Reporter Control |  | **** |
|  | AAV1 |  | **** |
|  | AAV2 |  | **** |
|  | AAV5 |  | **** |
|  | AAV8 |  | **** |
|  | AAV9 |  | **** |
|  | TransIT-X2 |  | **** |
|  | Lipofectamine 2000 |  | **** |
| TransIT-X2 | Reporter Control |  | **** |
|  | AAV1 |  | **** |
|  | AAV2 |  | **** |
|  | AAV5 |  | **** |
|  | AAV8 |  | **** |
|  | AAV9 |  | **** |
|  | Ad5/35 |  | **** |
|  | Lipofectamine 2000 |  | **** |
| Lipofectamine 2000 | Reporter Control |  | **** |
|  | AAV1 |  | ns |
|  | AAV2 |  | **** |
|  | AAV5 |  | **** |
|  | AAV8 |  | **** |
|  | AAV9 |  | **** |
|  | Ad5/35 |  | **** |
|  | TransIT-X2 |  | **** |

*Table S6. Cont.*
